# Native or promiscuous? Analyzing putative dimethylsulfoniopropionate lyases using a substrate proofing approach

**DOI:** 10.1101/168930

**Authors:** Lei Lei, Kesava Phaneendra Cherukuri, Diana Meltzer, Uria Alcolombri, Dan S. Tawfik

**Affiliations:** Department of Biomolecular Sciences, Weizmann Institute of Science, Rehovot 76100, Israel

**Keywords:** Enzyme promiscuity, enzyme superfamily, substrate profiling, dimethylsulfide

## Abstract

Enzyme promiscuity is widely spread. Foremost, within superfamilies, the native function of one enzyme is typically observed as promiscuous activity in related enzymes. The native function usually exhibits high catalytic efficiency while promiscuous activities are weak, but this is not always the case. Thus, for certain enzymes it remains questionable whether their currently known activity is native or promiscuous. Dimethylsulfon-iopropionate (DMSP) is an abundant marine metabolite cleaved via β-elimination to release dimethylsulfide (DMS). Eight different gene families have been identified as putative DMSP lyases, 5 of them belonging to the same superfamily (cupin-DLL; see the accompanying paper). Some of these enzymes exhibit very low activity, but this can be due to suboptimal folding or reaction conditions. We developed a substrate profiling approach with the aim of distinguishing native DMSP lyases from enzymes that promiscuously act as DMSP lyases. In a native DMSP lyase, relatively small changes in the structure of DMSP should induce significant activity drops. We thus profiled substrate selectivity by systematically modifying DMSP while retaining reactivity. Three enzymes that exhibit the highest activity with DMSP also exhibited high sensitivity to perturbation of DMSP’s structure (Alma, DddY, and DddL). The two enzymes with the weakest DMSP lyase activity also showed the highest crossreactivity (DddQ, DddP). Combined with other indications, it appears that the DMSP lyase activity of DddQ and DddP is promiscuous although their native function remains unknown. Systematic substrate profiling could help identify and assign potential DMSP lyases, and possibly applied to other enzymes.

**Abbreviations:** DMSP, dimethylsulfoniopropionate; DMS, dimethylsulfide; cupin-DLL, cupin DMSP lyase and lyase-like.

**Funding:** Financial support by the Estate of Mark Scher, and the Sasson & Marjorie Peress Philanthropic Fund, are gratefully acknowledged. D.S.T. is the Nella and Leon Benoziyo Professor of Biochemistry.

## Introduction

Enzymes are renowned for being able to convert one specific substrate into a given product with high turnover rates. Nonetheless, enzyme promiscuity, i.e., the ability to transform substrates other than the native substrate, or even to catalyze a different reaction, is widely spread.^*1*–*3*^ Take nitrogenase, to name just one example. Its native, physiological function is to reduce N_2_ to NH_3_. However, this enzyme reduces alternative substrates such as cyanide or acetylene,^*4*,*5*^ and the latter is widely applied as a surrogate of nitrogenase activity.^*6*^ Promiscuity is also a major driving force in the natural evolution of new enzyme functions.^*1*, *2*, *7*^

Within enzyme superfamilies, promiscuity reflects the catalytic mechanism shared by all superfamily members. A common key chemical step enables a range of reactions that can be catalyzed by the same configuration of catalytic active-site residues. For example, the enolase superfamily’s key chemical step is proton abstraction from a carbon adjacent to a carboxylate. This step underlies different reactions including dehydration, isomerisation or lactonisation, with a wide range of different substrates. ^*8*^ Owing to the shared chemistry, the native function of one superfamily member is often manifested as a promiscuous activity in related members and vice versa. This phenomenon helps unravel the evolutionary origins of enzymes,^*9*^ but has also led to mis-annotations.^*10*^ Take for example an enzyme called serum paraoxonase/ arylesterase, or PON1. This enzyme was characterized with synthetic substrates including paraoxon and phenyl acetate that are hydrolyzed with relatively high rates (*k_cat_*/*K_M_* ^~^ 10^4–^10^6^M^−1^s^−1^). However, PON1 turned out the be a lactonase, as indicated by systematically testing of different substrates, of which, only lactones exhibited a selectivity profile that is in agreement with a native function.^*11*^ Overall, although the native function typically occurs with high catalytic efficiency while promiscuous activities are weak, this criterion is not universally valid. In fact, the kinetic parameters for both native functions and promiscuous activities are distributed over a wide range.^*12*^ Thus, determining whether the known activity of an enzyme is native or promiscuous can be a challenge.

We encountered this challenge in the case of DMSP lyases. Many different genes encoding enzymes that catalyze the β-elimination of DMSP to release DMS a compound known as the ‘smell of the sea’, and acrylate are known (**Scheme**).

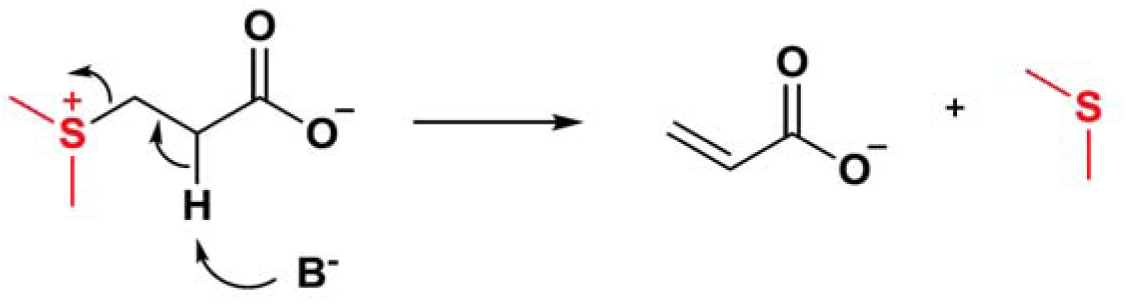

These putative DMSP lyases come from different families. Seven bacterial families, dubbed Ddd+, are known: DddD, a DMSP CoA-transferase-lyase (that generates 3-hydroxypropionyl-CoA rather than acrylate),^*13*, *14*^ DddP that belongs to the M24 proteinase family;^*15*^ and DddQ,^*16*^ DddL,^*17*^ DddW,^*18*^ DddK^*19*^ and DddY^*20*, *21*^ that all belong to a superfamily dubbed cupin-DLL as described in the accompany paper. One eukaryote family has also been discovered, Alma DMSP lyases, that belong to yet another superfamily (Asp/Glu racemase).^*14*^ Representatives from these families all show DMSP lyase activity, albeit at very different levels – the *k_cat_*/*K_M_* values range from < 1 M^−1^s^−1^ ^*16*, *22*–*24*^ up to 10^6^ M^−1^s^−1^.^*20*, *25*^ However, in itself, the low activity of some DMSP lyases such as DddQ and DddP is insufficient to indicate that DMSP lysis is merely a promiscuous activity of these enzymes. Low activity can also be the outcome of other factors, including mis-folding due to heterogeneous expression, incorrect cofactors such as metal ions, and/or suboptimal reaction conditions.

To distinguish enzymes whose native function is DMSP lyase from those in which this activity is likely to be promiscuous, we have developed an approach of substrate profiling. We have examined a series of DMSP analogues in which DMSP’s structure was systematically perturbed while retaining reactivity. We observed systematic behavior indicating high sensitivity to perturbation in some enzymes *versus* high cross-reactivity in others. Further, the specific activity of the 7 enzymes tested was found to be anticorrelated with their cross-reactivity. Overall, our results indicate that while DddL, DddY and Alma are *bona fide* DMSP lyases, the activity of DddQ and DddP is merely promiscuous.

## Results

### Substrate profiling of DMSP lyases

The rationale underlying substrate profiling is that, principle, if a given substrate is the enzyme’s native substrate, the enzyme’s active-site would have largely been shaped to fit this particular substrate. Thus, relatively small changes in the substrate’s structure are expected to induce relatively large drops in activity. True, the essence of enzyme promiscuity is acceptance of alternative substrates, sometimes with little resemblance to the original substrate and also in a selective manner.^*26*^ Nonetheless, selectivity is observed in most enzymes including promiscuous ones, and especially in enzymes that evolved for one particular substrate (in contrast to broad-specificity enzymes). As rules of thumb, substrates smaller than the original one will be accepted more readily than bigger ones (due to steric clashes) and increasingly larger substrates will show increasingly lower activity, and, perturbations next to where the key chemical step occurs would have larger impact.

In view of the above, a series of DMSP analogues was synthesized in which the potential for β-elimination and release of a dialkylsulfide product was retained (Figure 1A). The modifications included increasingly large dialkylsulfonyl leaving groups (EMSP, DESP, cDESP) and addition of methyl groups to the propionate moiety, either on the α-carbon from which a proton is abstracted (2-methyl-DMSP) or on the adjacent methylene (3-methyl-DMSP). None of these analogues matches a known natural metabolite except for DMSP of course.

**Figure 1.**
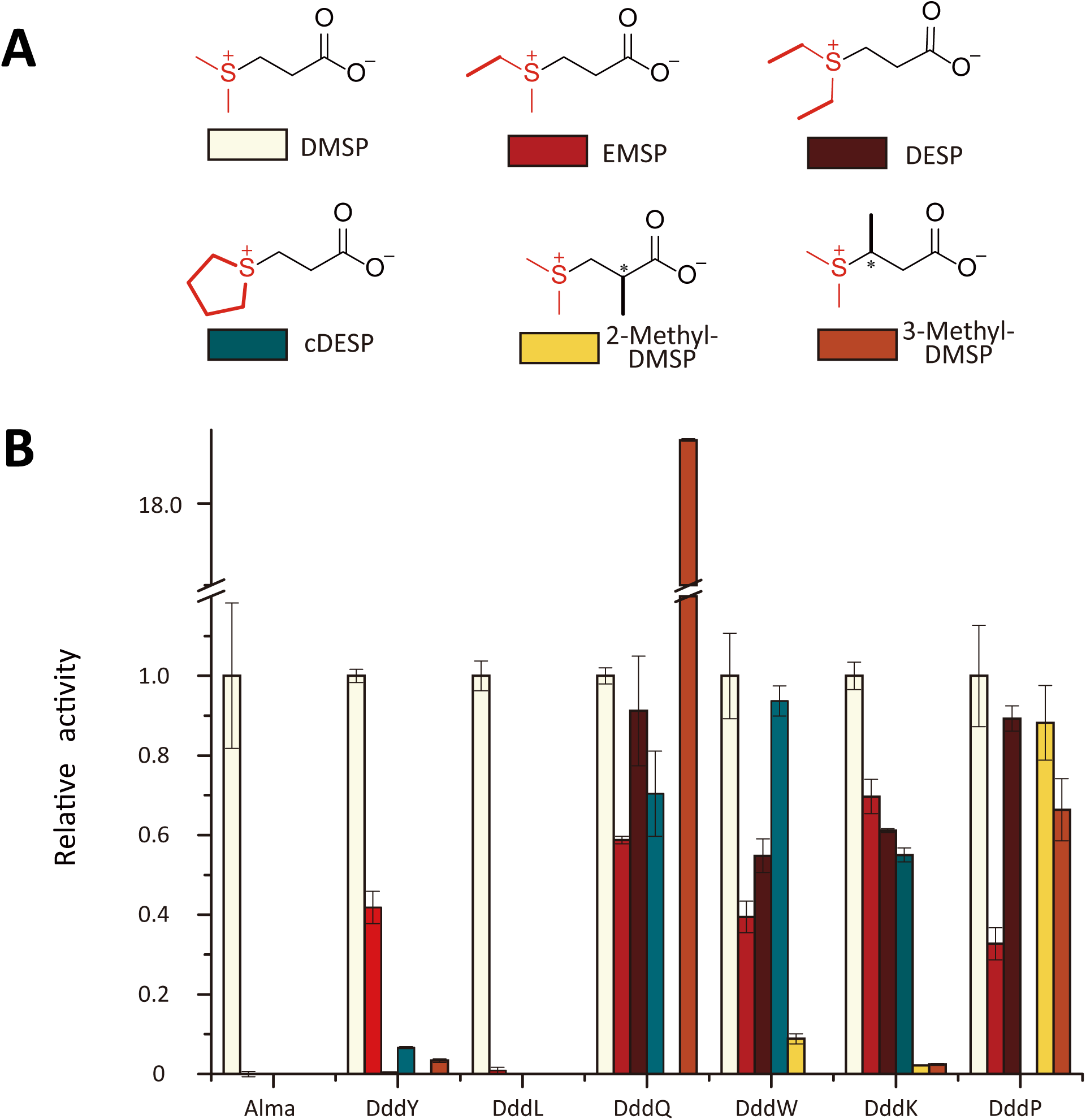
Profiling of DMSP lyases with a series of DMSP analogues. (A) The applied DMSP analogue. The dialkylsulfide leaving groups are denoted in red, and bold lines represent the modifications relative to DMSP. (B) The relative rates of elimination compared to the activity of these enzymes with DMSP. All substrates were applied at 10mM. Enzymes tested: Alma1 (*Emiliana huxleyi* Alma1,^27^ 0.3 μg/mL, DMS releasing rate, 114.5 μM/min); DddY (*Da*DddY, described in accompany paper, 20 ng/mL, DMS releasing rate, 27.8 μM/min); DddL^17^ (RsDddL; crude lysate, estimated enzyme concentration of 8 μg/mL (see accompanying paper), DMS releasing rate, 615.9 μM/min); DddQ^16^ (*Ruegeria pomeroyi* DSS-3, 100 μg/mL, DMS releasing rate, 13.2 μM/min); DddW^18^ (*Ruegeria pomeroyi* DSS-3, 15 μg/mL, DMS releasing rate, 119.9 μM/min); DddK^19^ (*Candidatus* Pelagibacter ubique HTCC1062, 8 μg/mL, DMS releasing rate, 121.92μM/min); DddP^22^ (*Roseovarius nubinhibens*, 10ug/mL, DMS releasing rate, 4.24μM/min).

We tested representatives of 6 bacterial Ddd+ families (all bacterial families except DddD which is a CoA-ligase/DMAP lyase confirmed as a *bona fide* DMSP lyase, ref^*14*^) and also the algal *E. huxeleyi* Alma1^*27*^. The bacterial DddY was expressed, purified and characterized as described in the accompanying paper. DddL could not be purified, but could be assayed in crude *E. coli* cell lysates (see also accompanying paper). DddK, DddW and DddP were cloned, expressed as purified as described in the Methods section. Specific activities and kinetic parameters are given in Table 1. These roughly match the reported ones although considerable differences were observed. For example, our preparation of *Ruegeria pomeroyi* DSS-3 DddQ exhibited >100-fold higher specific activity compared to a previous report.^*16*,*28*^ For DddK and DddW, although *K_M_* values have been reported,^*29*, *30*^ we observed no saturation with DMSP concentrations up to 10 mM and could therefore only measure *k_cat_*/*K_M_* (**Supplemental Figure S1**). Since all these bacterial enzymes are metal-dependent, activity differences may stem from different preparations containing different metals (but also from different assay conditions, *e.g.* buffer capacity, see Methods). Metal composition may have a different effect on the native compared to promiscuous activities, but these differences primarily regard catalytic promiscuity, *i.e.*, different reactions.^*31*^ Foremost, our substrate profiling approach regards the measurement of relative activities of DMSP analogues compared to DMSP, and hence the results should be relevant regardless of differences in specific activities. Indeed, the specific activities of the 7 tested enzymes with DMSP vary by >3,000-fold (Table 1). Thus, the enzymes were applied at different concentrations in accordance with their specific activities. The initial rates observed with each substrate analogue, for release DMS, or of the corresponding dialkyl sulfide, were compared to the initial rate of the same enzyme with DMSP (Figure 1B).

**Table 1:**
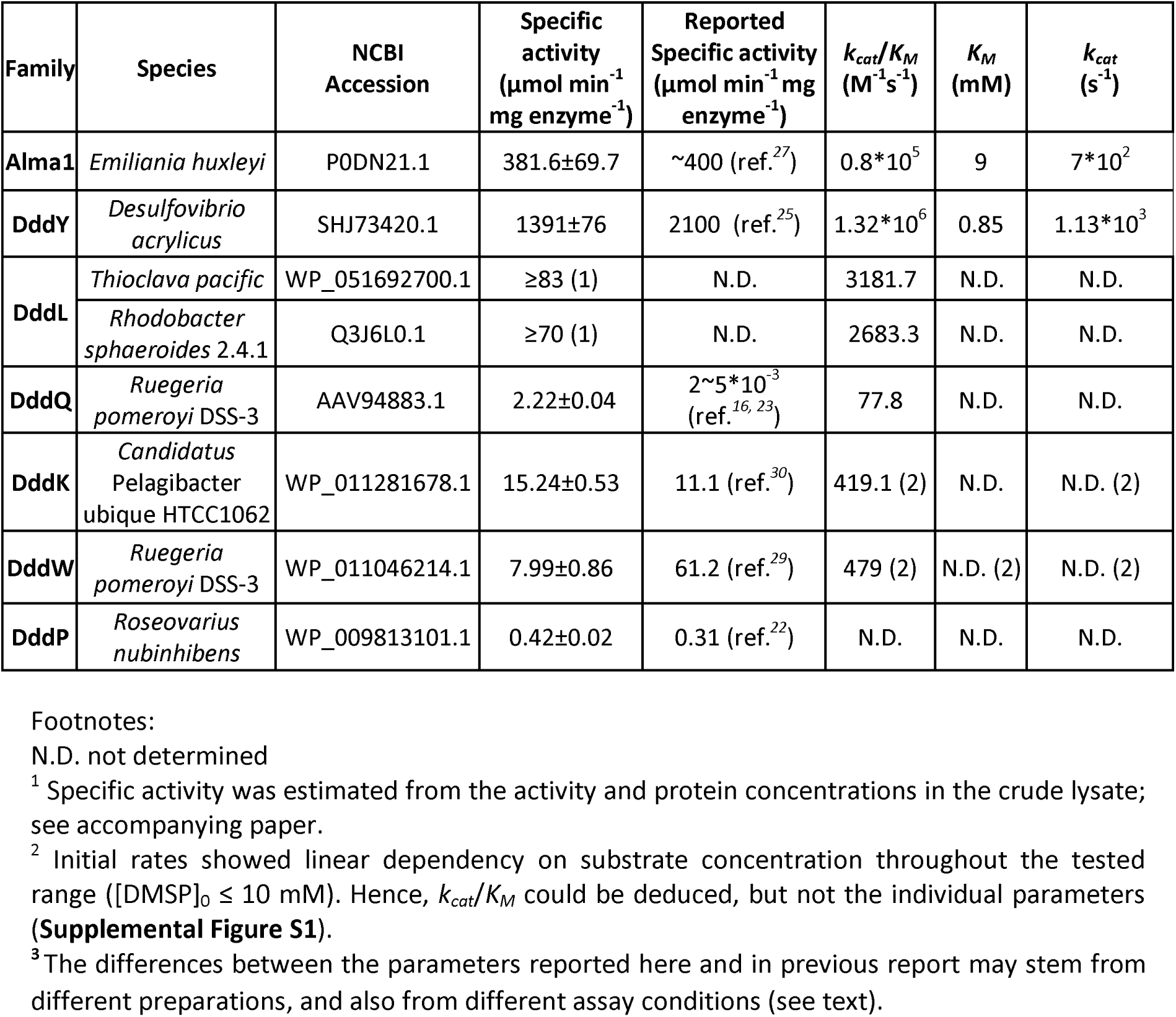
Kinetic parameters of the tested DMSP lyases.

### Alma1, DddLand DddY exhibit the expected DMSP-tailored selectivity

As can be seen in Figure 1B, the enzymes tested can be readily divided to two groups: highly selective enzymes (Alma1, DddY and DddL) and less, or evidently non-selective enzymes (DddK, DddW, DddQ, and DddP respectively). Specifically, Alma and DddL failed to accept any of the applied DMSP analogues under the tested conditions (enzyme concentrations and initial rates for DMS release with DMSP are noted in the legend). DddY showed some cross-reactivity. However, a significant and systematic decrease in activity was observed with increased size of substituents on the dialkylsulfide leaving group. Foremost, almost no activity was observed with analogues contacting an extra methyl group close to where proton abstraction occurs (2-methyl-DMSP and 3-methyl-DMSP).

Overall, profiling with DMSP supports the hypothesis that Alma1, DddY and DddL are *bona fide* DMSP lyases. This conclusion is also supported by their relatively high specific activity (elaborated below). However, in the other 4 bacterial Ddd+ families, low selectivity toward DMSP is evident.

### DddQ and DddP are highly likely to be promiscuous DMSP lyases

DddQ exhibited a trend that is in clear contrast to the selectivity observed in Alma1, DddY and DddL. Modifications of the leaving group, including the bulkiest one (cDESP) hardly affected the rate. Foremost, DddQ catalyzes the β-elimination of 3-methyl-DMSP nearly 20 times faster compared to DMSP. Considering that 3-methyl-DMSP is not a natural metabolite (as much as is known), DddQ is highly likely to be an enzyme that catalyzes a reaction other than DMSP lysis.

That DddQ is highly likely to be an enzyme whose native activity is not DMSP lyase is also supported by the genome neighborhood analysis presented in the accompanying paper. In bacterial genomes, enzymes that are functionally related are typically found in gene clusters, or even in the same operon. DddQ’s genome neighborhood does not seem to include acrylate catabolizing enzymes while genes encoding such enzymes are observed in DddD and DddY (see accompanying paper). In fact, the genome neighborhood suggests that DddQ is part of pathways(s) that catabolize proline-betaine or/and hydroxyproline-betaine. That 3-methyl-DMSP is a much better substrate for DddQ also suggests that DddQ’s real substrate is bulkier than DMSP.

Given the pattern of cross-reactivity, and its low activity, DddP is also likely to promiscuously act as DMSP lyase. The genome context of DddP is highly conserved (**Supplemental figure S2**). However, acrylate catalbolizing genes are not observed in DddP’s vicinity. The proximity of murein transglycosylase, type I glutamate—ammonia ligase, and a nitrogen regulatory protein, suggest that this gene cluster is involved in peptidoglycan recycling, cell division or nitrogen metabolism. However, what is the primary enzymatic function of DddP, or of DddQ, remains unknown.

## Discussion

Substrate specificity is never absolute, and many, if not most enzymes accept alternative substrates, promiscuously, or because they a priori evolved with broad substrate specificity.^*1*^ Identifying whether a given activity is promiscuous or native can be non-trivial for certain enzymes, and especially for enzymes in specialized metabolism. This challenge further intensifies in cases where specific activity is low, because low activity can also be the outcome of suboptimal expression or/and reaction conditions. In the case of DMSP lyases, proteins from 4 different superfamilies are known whose DMSP lyase activity spans from 0.4 up to 1,300 Units (Table 1). Nonetheless, there are clear indications for suboptimal folding, and also issues related to the incorporation of the catalytic metal in some of these enzymes; DddL, for example, could not be purified and could only be assayed in crude lyases (see accompanying paper). To circumvent these hurdles, we developed a substrate profiling approach.

### Quantifying substrate selectivity

Having obtained the substrate selectivity profiles of 7 different enzymes, we asked how correlated is their selectivity toward DMSP with their the level of their DMSP lyase activity. To examine such a correlation, a quantitative measure of enzyme selectivity had to be developed. Previous works provided quantitative measures of promiscuity, but these measures were designed to assess how broad is the substrate acceptance of a given enzyme. In contrast, we aimed at examining how likely is DMSP to be the native substrate. The more selective an enzyme is toward DMSP, the lower is the average relative activity with various DMSP analogues. In the simplest manner, a cross-reactivity index, *I_CR_*, can be defined as:

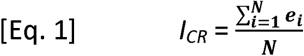

whereby *N* is the number of tested DMSP analogues, and *e_i_* is the activity relative to DMSP per each DMSP analogue. If an enzyme cleaves only DMSP, *I_CR_* =0, as is the case with Alma1 (Table 2). In principle, *I_CR_* is expected to be in the range of 0 up to 1. However, for DddQ the *I_CR_* value is >>1 due to the activity with one analogue being ^~^19-fold higher than with DMSP.

**Table 2:** Cross-reactivity of the tested DMSP lyases with DMSP analogues. The activities with the various analogues are normalized to the activity of the same enzyme

| Enzyme | EMSP | DESP | cDESP | 2-Methyl-DMSP | 3-Methyl-DMSP | $I_{CR}$ | $*I_{CR}$ |
| --- | --- | --- | --- | --- | --- | --- | --- |
| Alma1 | 0±0.006 | 0.00 | 0.00 | 0.00 | 0.00 | 0.00 | 0.00 |
| DddY | 0.42±0.04 | 0.0040±0.002 | 0.07±0.003 | 0.00 | 0.03±0.004 | 0.10 | 0.51 |
| DddL | 0.01±0.004 | 0.00 | 0.00 | 0.00 | 0.00 | 0.0017 | 0.008 |
| DddQ | 0.59±0.009 | 0.91±0.14 | 0.70±0.11 | 0.00 | 18.78±0.02 | 4.19 | 4.71 |
| DddW | 0.39±0.04 | 0.55±0.04 | 0.94±0.04 | 0.09±0.01 | 0.00 | 0.39 | 2.31 |
| DddK | 0.69±0.04 | 0.61±0.005 | 0.55±0.02 | 0.02±0.001 | 0.02±0.003 | 0.38 | 2.04 |
| DddP | 0.32±0.04 | 0.89±0.03 | 0.00 | 0.88±0.09 | 0.66±0.08 | 0.55 | 3.45 |
| STDEV | 0.25 | 0.39 | 0.37 | 0.30 | 6.53 |  |  |
| Mean | 0.35 | 0.42 | 0.32 | 0.14 | 2.79 |  |  |
| Coefficient of variation ($C_v^i$) | 0.71 | 0.91 | 1.14 | 2.14 | 2.35 | | |

with DMSP.

The above-described index has two drawbacks. Firstly, substrate analogues taken with rates much higher than the reference substrate shift certain enzymes drastically off scale (*e.g.* DddQ). Secondly, all analogues have the same weight. However, the applied substrate analogues structurally deviate from DMSP to different degrees – for example, cDESP presents a more dramatic structural perturbation than EMSP, and hence lack of cross-reactivity with the former is more indicative of a pocket shaped for DMSP. In principle, the more variable is the activity of different enzymes with a given analogue, the more relevant this analogue is as a reporter of the active-site’s complementarity to DMSP (in analogy to entropy, *H*, values applied in Ref. ^*32*^). To address the first issue, the activity was normalized per analogue (*i.e.*, dividing each activity (*e_i_*) by the highest activity observed with this analogue,
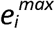). To address the second issue, for each analogue, the coefficient of variation, C_v_^i^, was calculated (the standard deviation for the normalized activities for all tested enzymes divided by the corresponding mean). The C_v_^i^ was applied to give each analogue a different weight. As can be seen in Table 1, the C_v_ values span within a reasonable range (0.7 – 2.4) and systematically reflect the structural deviations (gradually increasing from 0.7 to 1.14 with the bulkiness of the leaving group, and being with the highest for modifications close to the abstracted proton). The weighted cross-reactivity index ^*^*I_CR_*, is thus given by:

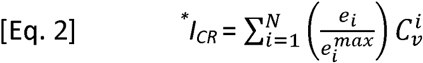

### Activity and selectivity correlate

Both cross-reactivity indexes seem to correlate with specific activity: the most active DMSP lyases are also the ones showing the clearest signature of a DMSP tailored active-site, and *vice versa* (Figure 2). The correlation is, as expected, not perfect. Some of the deviations may relate to suboptimal expression or/and reaction conditions; foremost, given the failure to purify DddL, this enzyme’s specific activity is most probably underestimated (see accompanying paper). Nonetheless, DddP and DddQ clearly stand out by both criteria – *i.e.*, they exhibit the lowest DMSP lyase activity as well as the lowest selectivity toward DMSP. This conclusion is in agreement with the genome context analysis (**Supplemental figure S2**; for DddQ see accompanying paper).

**Figure 2.**
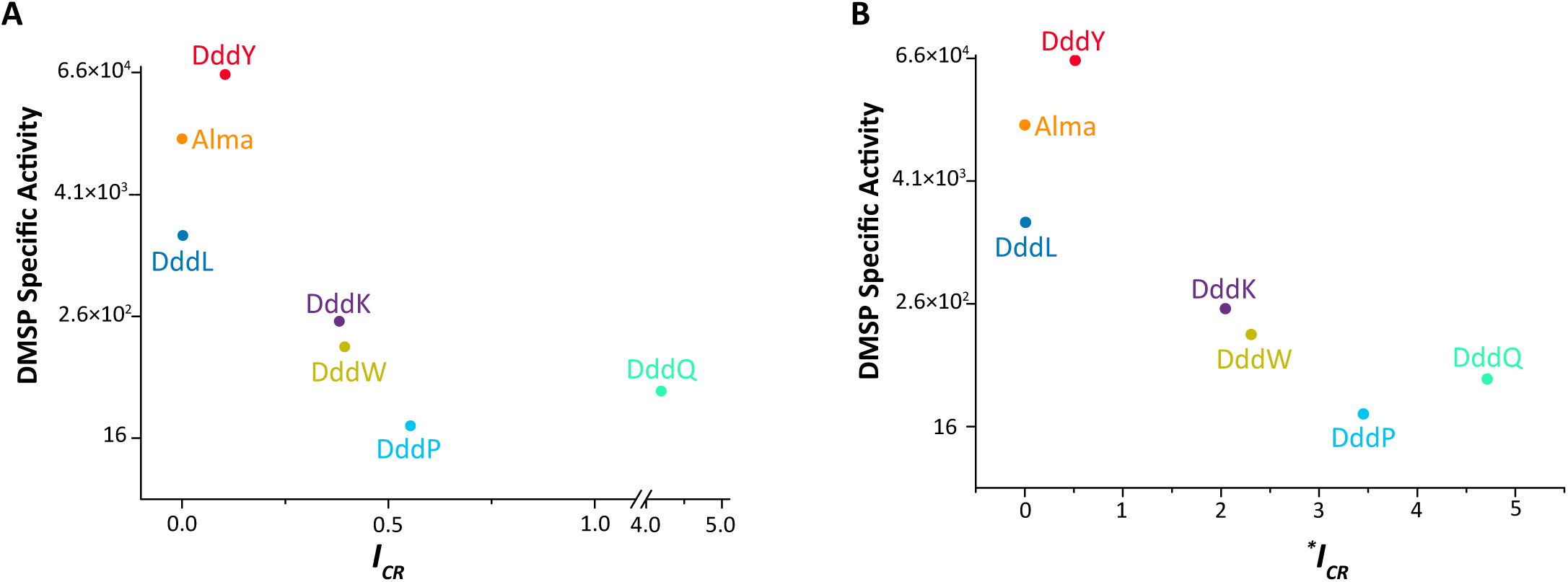
The DMSP lyase activity of the tested enzymes correlates with their selectivity toward DMSP. The Y-axis denotes the specific activity of the 7 enzymes tested (specified in Table 1) normalized by the enzyme’s molecular weight (the units are μmol DMS per min per μmol enzyme) and presented on a log2 scale. The x-axis denotes these enzymes’ cross-reactivity by two indexes: (A) *I_CR_* – the average relative activity with the 5 DMSP analogues (Equation 1); (B) ^*^*I_CR_* – a parameterized index of relative activities (Equation 2).

Overall, our analysis suggests that the DMSP lyases reported to date represent an entire range of evolutionary states. Some enzymes are fully diverged DMSP lyase specialists such as Alma, DddY and DddL. Other enzymes, most probably DddK and DddW, may comprise bi-functional intermediates – namely enzymes that possess the ancestral function (whatever this function might be) and have also evolved to have DMSP lyase activity. Bi-functional enzymes are common, especially in specialized metabolism.^*1*, *33*–*36*^ In other enzymes, DddQ and DddP in most likelihood, the DMSP lyase may be merely promiscuous, *i.e.*, a coincidental activity with no physiological relevance. Substrate profiling seems to reliably report these evolutionary states. Substrate profiling may be also valuable for mapping DMSP lyase activities of unknown origins (*i.e.*, by examining if they correspond to a highly selective DMSP lyase, or alternatively match the profile of one of the known promiscuous enzymes). Systematic substrate profiling might also be applicable to other enzyme classes in which it remains unclear whether the presently known activity in native or promiscuous.

## Material and Methods

### Enzyme cloning, expression and purification

The genes *dddK* and *dddW* were synthesized by Gen9. The synthesized gene fragments *dddK* and *dddW* were amplified and the PCR product was digested with *Nco*I and *Hind*III, and then cloned into the expression vector pET28a. This allowed DddK and DddW to be expressed with a C-terminal His-tag. Expression plasmid pET21a encoding *Roseovarius nubinhibens dddP*, and *Ruegeria pomeroyi* DSS-3 *dddQ*, were kindly provided by Professor Andrew Johnston, University of East Anglia. The recombinant enzymes were expressed in *E. Coli* BL21 (DE3). Cells were grown for overnight in 5 mL LB medium at 37°C. These cultures (1 mL) were used to inoculate 1 liter LB cultures that were subsequently grown at 37°C to OD_600nm_ of 0.6-0.8. The growth temperature was reduced to 16°C, and enzyme expression was induced with 0.1 mM IPTG. Following overnight growth at 16°C, the cells were harvested by centrifugation at 4°C. Cells were re-suspended in 50mL lysis buffer (100mM Tris-HCl pH 8.0, 100mM NaCl, 1mM CaCl_2_,10 mg lysozyme and 10 μg bezonase and lysed by sonication. Following centrifugation at 8,000×g for 20 mins, the clarified cell lysates were loaded on 2 mL Ni-NTA agarose beads (Millipore). Binding was performed at 4°C, and beads were washed with 50 mL lysis buffer followed by 100 mL lysis buffer with 35 mM imidazole. The enzymes were eluted with 150 mM imidazole. Fractions were analyzed by SDS-PAGE and assayed for DMSP lyase activity, combined, and the purified enzyme was concentrated by ultrafiltration (Amicon). Final enzyme concentrations were determined by the BCA assay. DddY, and DddL cloning, expression, and expression were described in accompanying paper, and Alma1 was expressed as described.^*27*,*37*^ To obtain the kinetic parameters, recombinant DddK, DddW and DddQ enzymes (at 8, 15 and 200 μg/mL respectively were reacted with DMSP at different concentrations up to 10mM for 5 mins (at these enzyme concentrations, rates of release were found to be linear up to 5 min). The amount of, the released DMS was measured to derive the initial rates (**Supplemental Figure S1**).

### DMSP lyase specific and cross activity assays

DMS release was measured as previously described.^*10*^ Briefly, freshly prepared 100 mM Tris-HCl pH 8.0 with 100 mM NaCl was used for the enzymatic assays supplemented with 10 mM DMSP as default. High buffer capacity is critical for assaying this reaction. Firstly, DMSP and its analogues are applied as hydrochloride salt as may just shift the pH. Additionally, every molecule of DMSP, or DMSP analogue, cleaved also releases one proton that affects pH. Reactions were performed at 30 °C (5 min, typically) and terminated by 1000-fold dilution into 30 mL chilled 10mM glycine pH 3.0 in sealed glass vials. Enzyme concentration were typically as follows: Ehux-Alma1, 0.3ug/mL; Sym-Alma1, 1μg/mL; *Desufovibrio* DddY 20ng/mL; *Alcaligenes* DddY 50ng/mL; DddW, 15μg/mL; DddQ, 100μg/mL; DddK, 8μg/mL. DddL was assayed in crude lysate, at an estimated concentration of 8ug/mL. DMS levels were determined using an Eclipse 4660 Purge-and-Trap Sample Concentrator system (OI Analytical) followed by separation and detection using GC-FPD (HP 5890) equipped with RT-XL sulfur column (Restek). All measurements were calibrated using DMS standards. Activity measurements of DMSP analogues were calibrated using the corresponding dialkylsulfides (ethyl methyl sulfite for EMSP, diethyl sulfite for DESP, and tetrahydrothiopene for cDESP).

### DMSP analogues synthesis

In general, all chemicals were purchased from Sigma-Aldrich and other commercial suppliers in reagent grade and used without further purification. Solvents are of AR grade and used without further purification. Deuterated solvents were from Sigma and Cambridge Isotope Laboratories, Inc. ^1^H and ^13^C NMR were recorded on BRUKER AVANCE III-400 (400 MHz) in D_2_O or CD_3_OD and all the signal positions were recorded in δ ppm with abbreviations s, d, t, q, m and dd denoting singlet, doublet, triplet, quartet, multiplet and doublet of doublet respectively. All NMR chemical shifts were referenced to residual solvent peaks (CD_3_OD δ =3.31 ppm and D_2_O δ =4.79 ppm). Coupling constants *J*, were registered in Hz. HRMS was determined with Xevo G2-XS QTOF mass spectrometer by electrospray ionization. The purity of all the compounds was more than 90% by ^1^H NMR.

The various DMSP analogues were synthesized by Michael addition of the corresponding dialkylsulfide and acrylic acid, or acrylic acid derivative, essentially as described.^*37*^ Typically, the corresponding acrylic acid (1 equivalent) was dissolved in 2M aqueous HCl (^~^10 mL for 1 gr), and the respective dialkylsulfide (3-6 molar equivalents) was added portion-wise. The reaction mixture was refluxed at 80°C for 3-12 hours. After cooling to room temperature, the solvent and the excess of unreacted dialkylsulfide were removed under reduced pressure. In the case of EMSP, DESP, cDESP, the resulting crude product (hydrochloride salt; solid, or syrup) was purified by recrystallization from isopropanol and in the case of 2-methyl DMSP and 3-methyl DMSP, the crude product hydrochloride salt; solid, or syrup) was triturated with diethyl ether. The products were dried under vacuum to obtain the hydrochloride salts. The products identity and purity were determined by ^1^H-NMR (400 MHz), ^13^C-NMR (101 MHz) and HRMS analysis (NMR spectra are provided in **Supplemental Figure S3**).

### DESP

obtained from diethyl sulfide and acrylic acid in 42% yield as a white solid.

^1^H NMR (400 MHz, D_2_O) δ 3.51 (t, *J* = 6.9 Hz, 2H), 3.38 (q, *J* = 7.4 Hz, 4H), 3.00 (t, *J* = 6.8 Hz, 2H), 1.47 (t, *J* = 7.4 Hz, 6H);^13^C NMR (101 MHz, D_2_O) δ 173.87, 33.48, 28.88, 7.89. HRMS *m/z* calcd for C_7_H_15_O_2_S^+^ is 163.0793, found 163.0801.

### EMSP

obtained from ethyl-methyl sulfide and acrylic acid in 44% yield as a colorless oil.

^1^H NMR (400 MHz, D_2_O) δ 3.62 – 3.55 (m, 1H), 3.52 – 3.45 (m, 1H), 3.45 – 3.38 (m, 1H), 3.38 – 3.29 (m, 1H), 3.01 (t, *J* = 6.8 Hz, 2H), 2.92 (s, 3H), 1.47 (t, *J* = 7.4 Hz, 3H); ^13^C NMR (101 MHz, D_2_O) δ 173.83, 36.49, 36.29, 28.66, 21.90, 7.79. HRMS *m/z* calcd for C_6_H_13_O_2_S^+^ is 149.0636, found 149.0641.

### cDESP

obtained from tetrahydrothiophene and acrylic acid in 20% yield as a white solid.

^1^H NMR (400 MHz, D_2_O) δ 3.69 – 3.57 (m, 2H), 3.55 – 3.47 (m, 2H), 3.44 (t, *J* = 6.8 Hz, 2H), 3.00 (t, *J* = 6.8 Hz, 2H), 2.44 – 2.24 (m, 4H);^13^C NMR (101 MHz, D_2_O) δ 174.01, 44.31, 37.99, 29.77, 28.22. HRMS *m/z* calcd for C_7_H_13_O_2_S^+^ is 161.0636, found 161.0630.

### 2-Methyl DMSP

obtained from DMS and 2-methylacrylic acid in a 15% yield as a white solid.

^1^H NMR (400 MHz, CD_3_OD) δ 3.57(dd, *J* = 13.3, 9.1 Hz, 1H), 3.44 (dd, *J* = 13.3, 5.1 Hz, 1H), 3.15 – 3.08 (m, 1H), 3.01 (s, 3H), 3.00 (s, 3H), 1.39 (d, *J* = 7.2 Hz, 3H); ^13^C NMR (101 MHz, CD_3_OD) δ 176.37, 48.09, 37.15, 27.28, 26.65, 17.13. HRMS *m/z* calcd for C_6_H_13_O_2_S^+^ is 149.0636, found 149.0640.

### 3-Methyl DMSP

obtained from DMS and trans-3-methylacrylic acid in a 10% yield as a white solid.

^1^H NMR (400 MHz, CD_3_OD) δ 3.96 – 3.85 (m, 1H), 3.01 – 2.91 (m, 8H), 1.57 (d, *J* = 6.9 Hz, 3H); ^13^C NMR (101 MHz, CD_3_OD) δ 172.65, 48.82, 36.87, 23.99, 22.35, 14.63. HRMS *m/z* calcd for C_6_H_13_O_2_S^+^is 149.0636, found 149.0643.

